# Development of conditional-siRNA programmable riboswitch for targeting adverse cardiac remodeling

**DOI:** 10.1101/2025.01.16.633434

**Authors:** Priyanka Gokulnath, Ane M. Salvador, Caleb Graham, Si-ping Han, Guoping Li, Ramaswamy Kannappan, Christopher Azzam, Michail Spanos, Lisa Scherer, Palaniappan Sethu, John Rossi, William A. Goddard, Saumya Das

**Author notes:** These authors contributed equally (Priyanka Gokulnath, Ane M. Salvador). **Address for correspondence:** Saumya Das, MD, PhD, Professor of Medicine, Cardiology Division, Cardiovascular Research Center, Massachusetts General Hospital, Harvard Medical School, Boston, MA 02114.

## Abstract

Heart Failure (HF) remains a global epidemic and a significant healthcare burden, with an unmet need for novel therapies to target the preceding pathological hypertrophy in vulnerable patients. Here we report the development of novel conditional-siRNA (*Cond-* siRNA) constructs that are selectively activated by disease-specific RNA biomarkers to enable cell-specific inhibition of a target disease-causing RNA. We designed a *Cond-* siRNA that can be activated by *nppa* mRNA, upregulated specifically in CMs under pathological stress, to silence the key pro-hypertrophic gene calcineurin by the effector siRNA. In cellular models including neonatal rat ventricular myocyte (NRVM) and rat cardiomyocyte cell-line (H9C2), *Cond-*siRNA exhibited low baseline activity in the absence of the disease biomarker but achieved targeted calcineurin silencing upon *nppa* mRNA induction by phenylephrine (PE)-induced stress in a two-dimensional (2D) cell culture system and pressure overload in three-dimensional (3D) heart-on a chip system. NRVM transfection with the *Cond-*siRNA resulted in a decreased expression of calcineurin mRNA specifically after PE or pressure-overload treatment, but not after vehicle treatment, proving *nppa* mRNA-specific activation of the effector siRNA against calcineurin. Specificity was confirmed as *Cond-*siRNA did not significantly silence calcineurin in cardiac fibroblasts and T cells, lacking *nppa* expression. Reduced calcineurin protein levels and NFATc1 nuclear translocation correlated with decreased NRVM hypertrophy after PE treatment, confirming *Cond-*siRNA’s efficacy. This study offers proof-of-concept for *Cond-*siRNA as a targeted therapy to mitigate hypertrophic progression, paving the way for novel HF treatments.

**One sentence summary:** Conditional-siRNA targeting adverse cardiac remodeling

**GRAPHICAL ABSTRACT:** 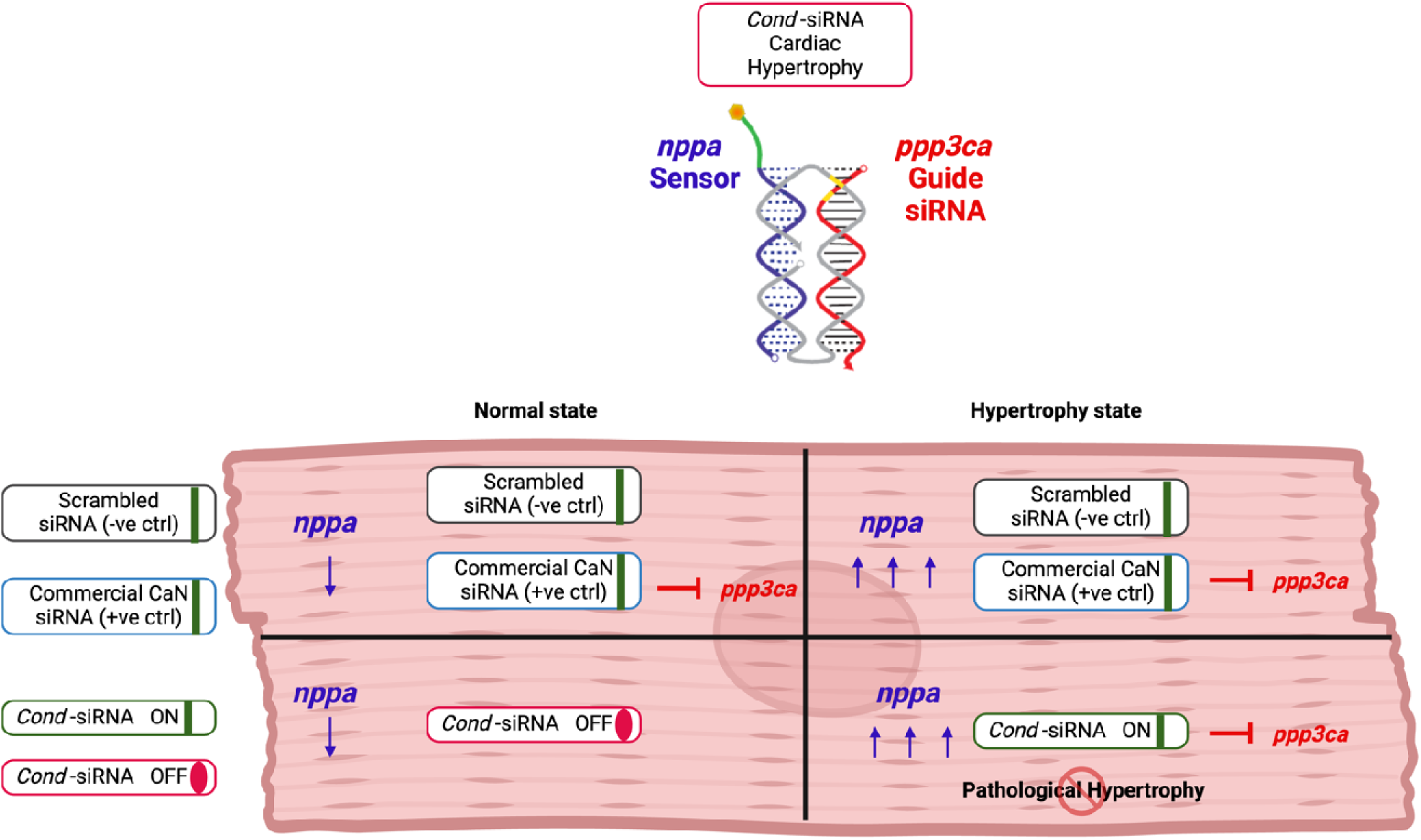

## INTRODUCTION

Heart Failure (HF) is a leading cause of morbidity and mortality worldwide, affecting over 26 million people worldwide, including 5.7 million in the United States, and represents a major contributor to healthcare expenditure in the US.^1,2^ Despite advances in HF treatment to reduce mortality, over 50% of patients die within 5 years from initial diagnosis.^3^ The global burden of HF continues to escalate, with a projected 46% increase in HF prevalence by 2030.^3^ According to the latest American Heart Association (AHA) / European Society of Cardiology (ESC) Heart Failure Guidelines,^4,5^ current therapeutic strategies primarily focus on symptom management and delaying disease progression, yet significant challenges remain in preventing the transition from compensatory hypertrophy to decompensated HF. These alarming trends of the HF epidemic underscore an unmet clinical need for effective therapies that can mitigate HF progression from its preceding pathological hypertrophy in at-risk patients.

Myocardial hypertrophy, an early compensatory response of the heart to pathological stress, is characterized by an upregulation of a specific transcriptional program.^6^ Over time, this adaptive response of cardiac hypertrophy progresses to cardiac decompensation and eventually clinical HF. ^7^ Experimental studies in animal models have identified genes that are critical mediators of this adverse cardiac hypertrophy that precedes HF.^8^ Notably, genetic and pharmacologic inhibition of key molecules that regulate CM hypertrophy pathways, such as calcineurin (CaN), have shown promise in blocking pathological hypertrophy and its progression to HF.^9–11^ However, translating these findings into clinical practice remains challenging due to the off-target effects in other non-cardiac cell types. For example, the critical role of CaN in immune cells suggests that its inhibition in a non-specific manner may lead to adverse side effects.^12^

Recent advances in RNA biology have highlighted its potential for serving as effective therapeutic targets and prompted the development of novel conditional-siRNA (*Cond-*siRNA constructs) or riboswitches.^13^ These molecules are specifically activated by signature RNAs that have low baseline expression in normal cellular conditions but are markedly increased in disease states, thereby serving as highly specific markers of pathology in the stressed cell. This enables the cell-specific and state-specific inhibition of a target RNA of the effector site of the *Cond-*siRNA construct. Such a conditional activation confers a high degree of specificity, enabling targeted silencing of disease-associated RNAs and minimizing off-target effects.

In this study, we hypothesize that a riboswitch-based *Cond-*siRNA will ablate pro-hypertrophic signaling, specifically in hypertrophied cardiomyocytes (CMs), and thus ameliorate HF progression. Specifically, we describe an RNA riboswitch that is activated by an mRNA transcript uniquely expressed in hypertrophied CMs, enabling targeted silencing of the calcineurin gene—a critical mediator of pathological hypertrophy and a novel strategy for HF treatment.

Expanding on our prior study where we demonstrated the feasibility of designing a *Cond*-siRNA to function as prescribed, we report here on the design and experimental validation of a *Cond-*siRNA specifically targeting cardiac hypertrophy in several models. We identify specific molecular sensors for riboswitch activation, evaluate their functionality, and demonstrate their efficacy in validated cardiac hypertrophy cell culture models as well as in a tissue-on-chip model. Our findings reveal that this riboswitch-based *Cond-*siRNA effectively inhibits key pathways involved in adverse cardiac remodeling, achieving robust knockdown activity with minimal off-target effects. These results, in line with our previous proof-of-principle evidence, offer a promising strategy for the targeted treatment of HF, representing a significant advancement in RNA-based therapeutics.

## RESULTS

### Identification of specific sensors and design of the *Cond-*siRNA targeting cardiac hypertrophy

To identify mRNA transcripts that would serve as sensors suitable to activate the *Cond-*siRNA, we screened several potential sensors by subjecting Neonatal Rat Ventricular Myocytes (NRVMs) to two stress models: **(1)** 24 hours of hypoxia (0.2% O_2_) followed by 12 hours of reoxygenation (Supplementary Figure 1A), and **(2)** treatment with the pro-hypertrophic molecule phenylephrine (PE) (Supplementary Figure 1B). These models mimic pathological events that occur in the myocardium during adverse cardiac remodeling. We extended this potential sensor mRNAs’ screen to another *in vivo* model by examining their expression in heart tissues from mice subjected to either non-ischemic (Transverse Aortic Constriction -TAC-) or ischemic (ischemia-reperfusion -IR-) HF (Supplementary Figure 1C). These experiments led to the selection of *nppa*, which encodes for ANP (Atrial Natriuretic Peptide), as the sensor of the *Cond-*siRNAs, based on its consistent upregulation in all *in vitro* and *in vivo* cardiac stress conditions.

### Synthesis of a Conditional-siRNA (*Cond-*siRNA) Construct Based on RNA Sensor *nppa* and Target Gene *Calcineurin*

The design of the Conditional-siRNA (*Cond-*siRNA) constructs paired the *nppa* RNA sensor with a target sequence for *calcineurin* (PPP3CA), using a systematic and iterative protocol to ensure specificity and functionality. The process began with the selection of a validated siRNA sequence targeting *calcineurin,* which was used as the guide strand. To create a functional 23-base pair Dicer substrate, four G/C-rich bases were added to the 5’ end of the guide strand, enhancing its stability and processing efficiency. For the input biomarker strand, the sequence was derived from the 3’ untranslated region (3’UTR) of *nppa* mRNA, a region known for its consistent upregulation during cardiac stress. Computational screening identified potential 31–33 nucleotide antisense segments, with candidates prioritized based on ∼50% GC content and the absence of problematic motifs such as GGGG or MMMM (M=A/U). The reverse complement of each segment was generated as the candidate sensor sequence, which was then validated using NCBI BLAST to exclude unintended matches to the mouse or rat genome.

Using the selected sensor strand, complementary segments of 11 and 12 bases were generated as the putative core strand, designed to interact with the first 23 bases of the sensor strand. Thermodynamic properties, such as binding affinity and secondary structure, were evaluated using Nupack, a nucleic acid ensemble structure prediction package.^14^ This analysis confirmed that the sensor strand exhibited minimal secondary structure and predicted strong and accurate binding between the sensor strand and the core strand’s 5’ and 3’ overhangs (Supplementary Figures 2A–B). The core strand was then finalized according to the complementary pattern for the *Cond-*siRNA design. For example, the sequences for the guide strand (CG AG UGUUGU UUGGC UU UUCCUG UU), the sensor strand (CUUCACCACCU CUCAGUGGCAAU GCGACCAA), and the core strand (AGGUGGUGAAG CAGGAAAAGCCAAACAACACUCGAUUGCCACUGAG) were systematically aligned to achieve precise duplex formation. Further analysis using Nupack confirmed minimal secondary structure in the core strand overhangs and ensured reliable binding between all components (Supplementary Figure 2C).

Chemical modifications were added to the *Cond-*siRNA strands to optimize functionality and stability. These modifications included Locked Nucleic Acid (LNA) substitutions for enhanced target binding affinity and conjugation with cholesterol to improve cellular uptake. An improved version of the first-generation construct (1^st^ G) had reduced phosphorothioate modifications of the sensor and an additional LNA modification in the middle toehold. Based on this 1^st^ G construct, a second generation (2^nd^ G) construct was altered with additional 2’ O methyl modifications of the guide strand to increase stability and reduce potential off-target effects. Further refinements were made to improve delivery by conjugating the new sensor with cholesterol (Chol. Conj) and triethylene glycol (Figure 1A). Following synthesis, the *Cond-*siRNA strands were mixed in equimolar concentrations in PBS buffer and thermally annealed using a thermocycler to facilitate precise self-assembly through base-pairing. The assembled *Cond-*siRNAs were then purified via native gel electrophoresis, which confirmed successful assembly, as evidenced by a clean band corresponding to the *Cond-*siRNA construct (Figures 1B and 1C).

**Figure 1.**
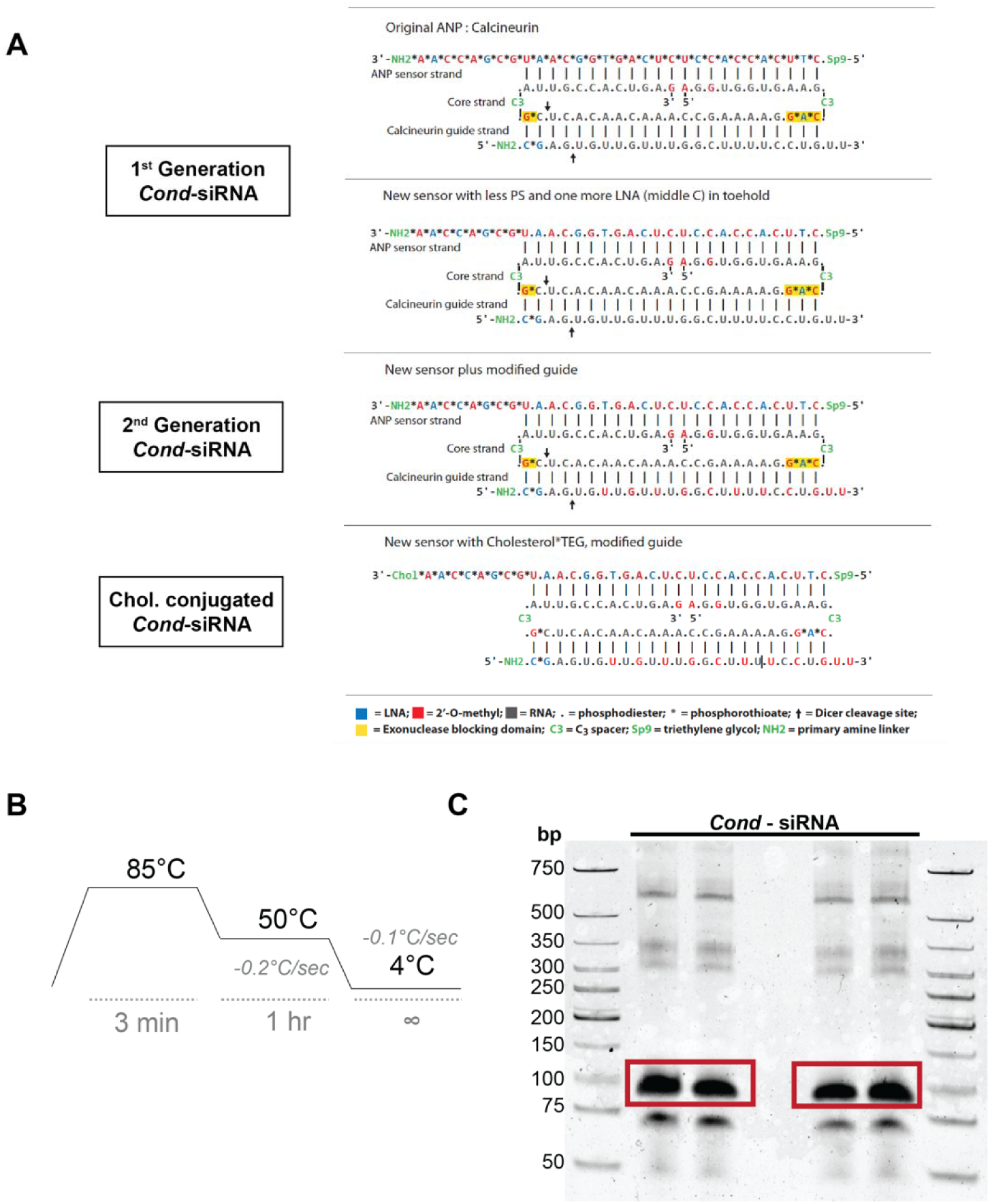
Synthesis of a Conditional-siRNA (*Cond*-siRNA) construct based on RNA sensor *nppa* and target gene calcineurin. **A.** *Cond*-siRNA constructs’ designs, based on *nppa* mRNA sensor and the target gene calcineurin. Each of the different constructs bears different chemical modifications, indicated below color-coded, which confer thermodynamic stability or cell-penetrating capacity. **B.** Thermocycler program used to induce base-pairing driven self-assembly of the different strands combined in an equimolecular manner to generate the *Cond*-siRNA via thermal annealing. **C.** Assembled *Cond*-siRNA constructs run in a 10% TBE appear as a clean band (highlighted in a red square). Higher molecular weight products correspond to concatemers of the different RNA strands, and smaller molecular weight products correspond to strands that did not assemble in the *Cond*-siRNA constructs.

This approach represents the successful design of a *Cond-*siRNA targeting *calcineurin*, a key gene involved in pathological cardiac hypertrophy. The method combines advanced computational tools, biochemical analysis, and chemical optimization to ensure high specificity, efficient assembly, and effective cellular delivery. This novel strategy lays the groundwork for therapeutic applications targeting adverse cardiac remodeling with precision and minimal off-target effects.

The annealing and purification process for Cond-siRNA constructs was optimized to produce well-assembled, concentrated constructs suitable for high-dose transfections. Strand annealing was tested under varying temperatures and salt concentrations to maximize yield while minimizing concatemers and unannealed strands. As shown in Supplementary Figure 3A, optimal results were achieved using 1× PBS as the annealing buffer, avoiding the high-molecular-weight concatemers observed at low (0.1× PBS) and high (5–10× PBS) salt concentrations. Post-purification, Cond-siRNA concentrations were increased from 2 µM to approximately 6 µM using ammonium citrate and ethanol-based methods (Supplementary Figure 3A). To further ensure proper assembly and conformation, a "re-annealing" step at 50°C was introduced (Supplementary Figure 3B), resulting in high-quality Cond-siRNA constructs free of by-products. This optimized methodology provides well-concentrated Cond-siRNA constructs for targeting key mediators of pathological cardiac hypertrophy.

### Evaluation of the *in vitro* efficiency of activation of the *Cond-*siRNA construct by the sensor and ability to block its target gene

To assess the efficiency of *Cond-*siRNA activation and its ability to block *calcineurin* expression, we evaluated its uptake, activation, and specificity in *in vitro* models. NRVMs were transfected with FITC-labeled *Cond-*siRNA constructs, either unconjugated or cholesterol-conjugated, to visualize cellular uptake. Immunofluorescence imaging confirmed successful uptake, with FITC signals colocalizing with cardiomyocyte-specific troponin staining (Figure 2A). While RNAiMax was required for the transfection of unconjugated *Cond-*siRNA, cholesterol-conjugated constructs entered cells without the need for transfection reagents, highlighting their enhanced delivery potential.

**Figure 2.**
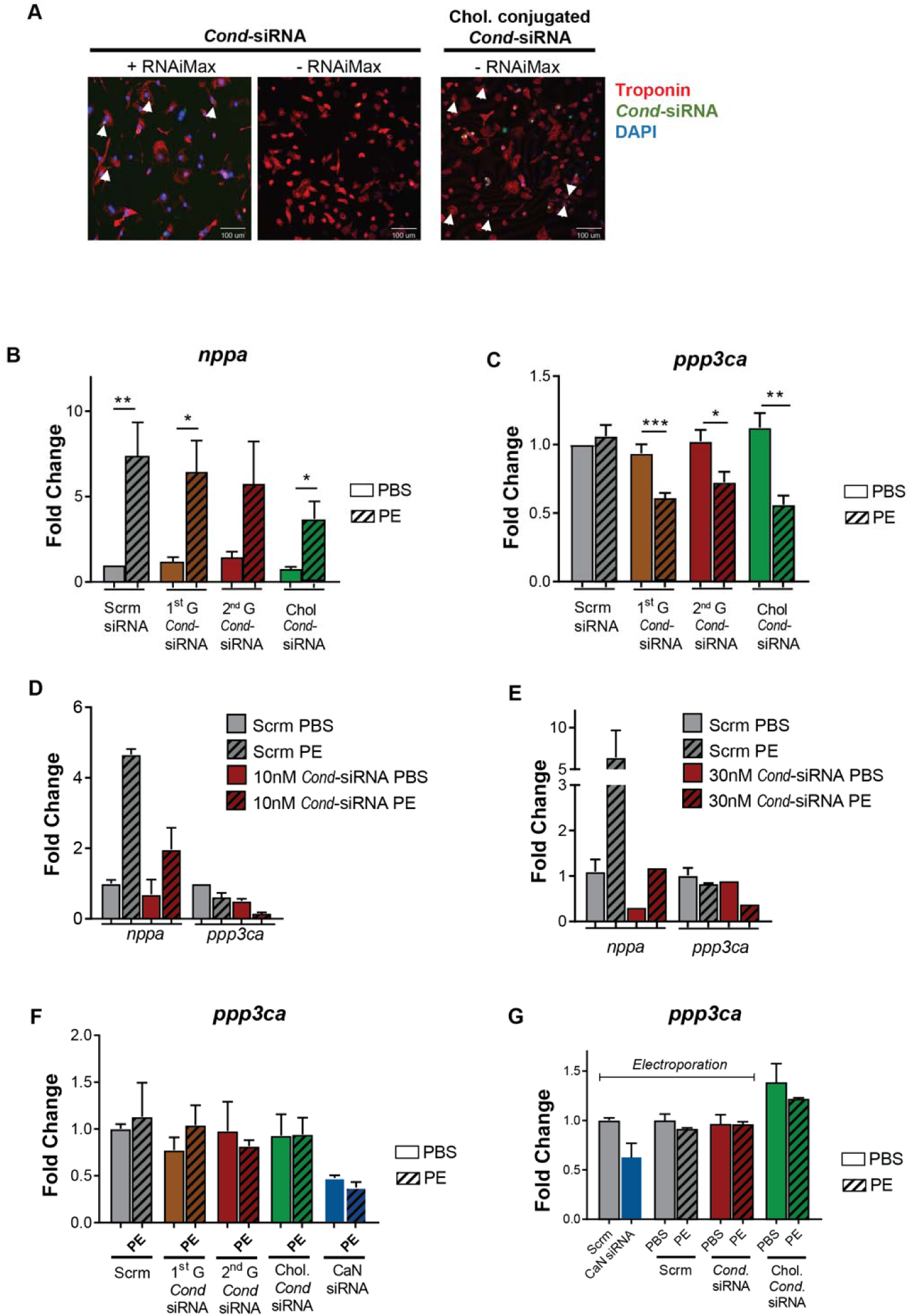
Cardiomyocyte specific activation and siRNA activity of *Cond*-siRNA. **A.** Successful transfection of *Cond*-siRNA with lipofectamine RNAiMax or without any transfection reagent in cholesterol conjugated *Cond*-siRNA into NRVM. *Cond*-siRNA is labeled with FITC, and NRVMs are stained with Troponin T antibody (red) and DAPi nuclear staining (blue). **B-E.** Calcineurin (*ppp3ca*) and ANP (*nppa)* mRNA expression in NRVMs treated with either 1nM (**B-C**) of the different *Cond*-siRNA constructs or **D**. 10 and **E.** 30nM of *Cond*-siRNA. 24h post-isolation, NRVMs were transfected with the *Cond*-siRNA; after 24h, NRVMs were treated with 50µM PE for 48h, and RNA expression was determined 72h post-transfection. Data is shown as fold vs scramble (Scrm) PBS. N=4-9. Unpaired t-test was performed between PBS and PE groups, with significance indicated as *p 0.05, **p 0.01, ***p 0.001. **F.** Calcineurin (*ppp3ca*) mRNA expression in NRCFs treated with 1nM of the different *Cond*-siRNA constructs or the commercial calcineurin siRNA. 1-week post-isolation, NRCFs were transfected with the *Cond*-siRNA; after 24h, NRCFs were treated with 50µM PE for 48h, and RNA expression was determined 72h post-transfection. Data is shown as fold vs scramble (scrm) PBS. N=3. **G.** Calcineurin (*ppp3ca*) mRNA expression in Jurkat T cells electroporated or treated with 1nM of commercial siRNA (CaN siRNA) or *Cond*-siRNA constructs. 24h after treatment with the *Cond*-siRNA cells were treated with 50µM PE for 48h, and RNA expression was determined 72h post-treatment. Data is shown as fold vs scramble (scrm) PBS.

Having demonstrated the uptake of the *Cond-*siRNA, the activation and siRNA activity efficacy of the *Cond-*siRNA was evaluated in NRVMs under PE-induced stress conditions (Supplementary Figure 4A), as compared to the vehicle. NRVM transfection with 1nM of first-generation (1^st^ G), second-generation (2^nd^ G) or cholesterol-conjugated (Chol) *Cond-*siRNA construct resulted in a 50% decrease in the expression of calcineurin only after PE treatment (Figure 2B). Importantly, no reduction in *calcineurin* expression was observed in vehicle-treated cells, further validating the conditional nature of the constructs, which respond specifically to PE-induced *nppa* upregulation (Figure 2C). Transfection of NRVMs with higher doses (10 nM and 30 nM) of the *Cond-* siRNA (Figure 2D-E) led to a further reduction in calcineurin silencing, achieving a 70-80% decrease in expression. The degree of calcineurin silencing achieved with the *Cond-*siRNA was similar to the one observed when transfecting commercial calcineurin targeting siRNA into NRVMs (Supplementary Figure 4B).

With the goal of demonstrating the cardiomyocyte-specific activation and siRNA activity of the *Cond-*siRNA, Neonatal Rat Cardiac Fibroblast (NRCF) was utilized to quantify the expression of calcineurin after treatment with 1 nM *Cond-*siRNA (2^nd^ G) in the presence of PE (Figure 2F). As expected, the *Cond-*siRNA did not reduce calcineurin expression in the presence of PE, reaffirming the cell-specific activation and siRNA activity of the *Cond-*siRNA, restrained to *nppa* expressing NRVMs. Furthermore, considering the important function of calcineurin in T cell activation and effector function, the *Cond-*siRNA activity was determined in immortalized human Jurkat T cells (Figure 2G, Supplementary Figure 5). Given that the *Cond-*siRNA gets activated only in the presence of *nppa* mRNA, the construct should not get activated in T cells (see *nppa* amplification plot in Supplementary Figure 5), and therefore, it would not alter T cell activation and immune competence. To test this, siRNA transfection conditions were optimized through electroporation (Supplementary Figure 5). While electroporation of Jurkat cells with a commercial siRNA targeting calcineurin led to ∼50% decrease in calcineurin expression, treatment with 1nM *Cond-*siRNA and cholesterol-conjugated *Cond-*siRNA did not result in significant changes in calcineurin expression in the presence or absence of PE, further validating cardiomyocyte *nppa* specific activation of the constructs and therefore reassuring lack of immune-deficient effect when using the *Cond-*siRNA *in vivo*.

### Treatment of NRVMs with the *Cond-*siRNA leads to inhibition of NFATc1’s nuclear translocation and pathological hypertrophy without promoting myocyte atrophy

Having demonstrated the siRNA activity of the *Cond-*siRNA in NRVMs in the presence of PE-induced upregulation of ANP, a characterization of calcineurin downstream signaling based on prior research^15,16^ and the expression of pathological cardiac hypertrophy markers was performed in the presence of the *Cond-*siRNA and PE. In line with RNA expression, protein levels of calcineurin in NRVMs were found to be reduced in the cytoplasm after treatment with 1 nM *Cond-*siRNA in the presence of PE (Figure 3A), resulting in a decreased translocation of NFATc1 into the nucleus (Figure 3B). Interestingly, when quantifying the expression of pathological myosin heavy chain β isoform protein levels, a lack of upregulation and overall decreased expression of the isoform was observed in the presence of the *Cond-*siRNA and PE (Figure 3C). Furthermore, phosphorylation of ERK1/2 was also decreased in NRVMs treated with the *Cond-*siRNA in response to PE, as compared to Scramble siRNA-treated cells (Figure 3C). Together, these data suggest a potent effect on pathways related to pathological cardiac hypertrophy with Cond-siRNA in a state-specific manner.

**Figure 3.**
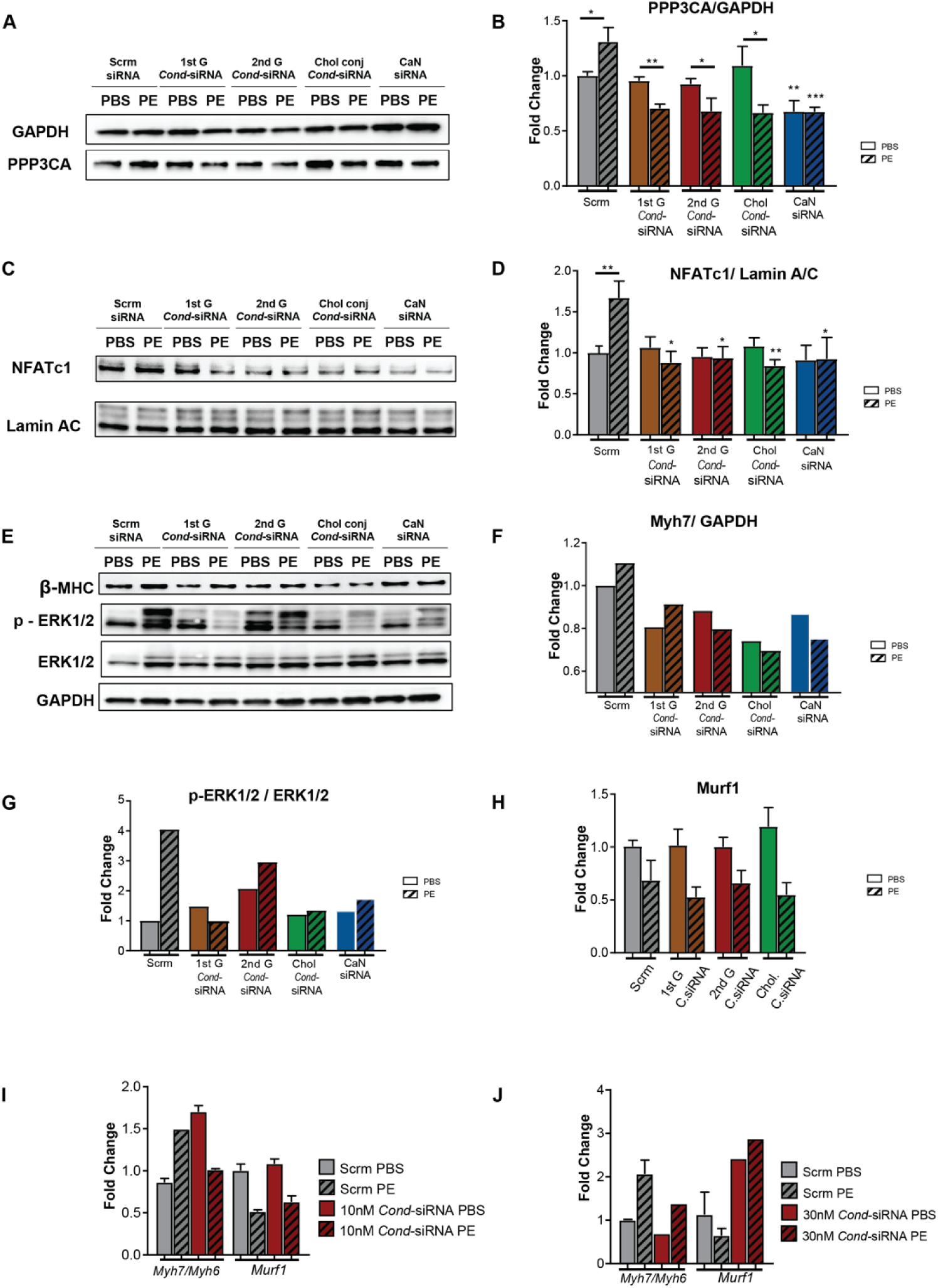
Inhibition of NFATc1’s nuclear translocation and pathological hypertrophy without promoting myocyte atrophy. **A-G.** Quantification of protein levels of **A-B.** calcineurin in the cytoplasm, **C-D**. NFATc1 in the nucleus, **E-G.** -MHC and p-ERK1/2/ERK1/2 in NRVMs treated with 1 nM of the commercial siRNA (CaN siRNA) or the different *Cond*-siRNA constructs. 24h post-isolation, NRVMs were transfected with the *Cond*-siRNA; after 24h, NRVMs were treated with 50µM PE for 48h, and protein expression was determined 72h post-transfection. Data is shown as fold vs scramble (scrm) PBS. Unpaired t-test was performed between PBS and PE groups, with significance indicated as *p 0.05, **p 0.01, ***p 0.001. **H-J.** Quantification of mRNA levels of atrophy marker H-J. *Murf1* and **I-J.** MYH7/MYH6 in NRVMs treated with 1nM (**H**), 10nM (**I**), and 30nM (**J**) of the different *Cond*-siRNA, treated without 50µM PE for 48h. Data is represented as fold vs Scramble PBS. n=4. Statistics One-Way ANOVA vs Scramble PBS.

In order to evaluate the potent anti-hypertrophic effects of the Cond-siRNA may, on the other hand, upregulate pathways of muscle atrophy (which would be deleterious as a treatment side-effect). To test this, mRNA levels of *Murf1* were quantified, finding that no significant upregulation was observed at 1nM and 10nM concentration (Figure 3D). This mimics the expression of this atrophy marker in NRVMs treated with commercial siRNAs targeting calcineurin (Supplementary Figure 6A). Interestingly, further silencing of calcineurin using higher concentrations of *Cond-*siRNA (30 nM) or commercial siRNA (50 nM and 100 nM, Supplementary Figure 6A) led to an increase in the expression of atrophy marker *Murf1*. As a positive control of muscle atrophy, dexamethasone and miRNA-29b were utilized,^17^ where its treatment or transfection to NRVM led to an upregulation of the atrophy marker *Murf1* (Supplementary Figure 6B). Therefore, the upregulation of atrophic genes when severely silencing calcineurin expression, points towards a therapeutic window to decrease calcineurin levels but without inducing NRVM atrophy.

Calcineurin silencing mediated by the *Cond-*siRNA prevents PE-induced cardiomyocyte hypertrophy. With the goal of assessing the corresponding phenotypical effect of calcineurin silencing by the *Cond-*siRNA on the cells, the NRVM area was quantified in the presence of the different generations of the *Cond-*siRNA, in the presence or absence of PE. Staining of NRVMs with troponin enabled the tracing and measurement of cell area and showed that the reduced calcineurin expression in cells treated with the *Cond-*siRNA exposed to PE resulted in a reduced cellular hypertrophy phenotype, presenting significantly lower NRVM area after PE treatment, as compared to scramble siRNA treated NRVMs (Figure 4).

**Figure 4.**
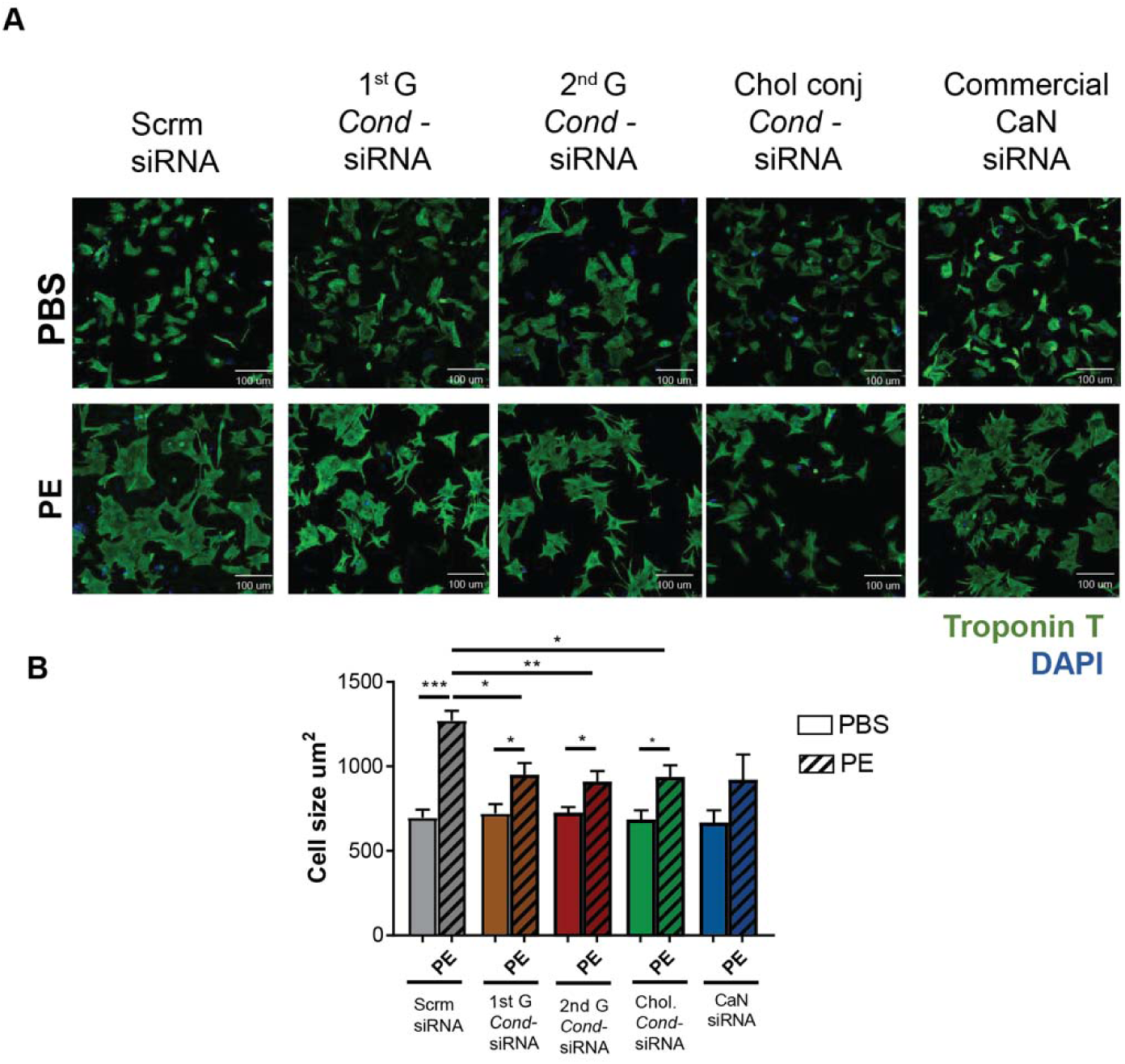
**A.** Representative images and **B.** Quantification of NRVM area as a marker of cellular hypertrophy in NRVMs treated with 1nM of the different *Cond*-siRNA constructs or the commercial calcineurin (CaN) siRNA 24h post-isolation, NRVMs were transfected with the *Cond*-siRNA; after 24h, NRVMs were treated with 50µM PE for 48h and cellular area was measured 72h post-transfection having stained the cells for Troponin T. N=3-5. Unpaired t-test was performed between PBS and PE groups, with significance indicated as *p 0.05, **p 0.01, ***p 0.001.

Altogether, we have characterized in depth the cardiac myocyte remodeling phenotype in response to the PE and *Cond-*siRNA treatment, proving target silencing and phenotypical response only in cells expressing the *Cond-*siRNA activating sensor *nppa*.

### Evaluation of *Cond*-siRNA in a Heart-On-Chip (HOC) Model under pressure-overload condition

A preclinical Heart-On-Chip (HOC) model was successfully developed previously to simulate HF conditions induced by pressure overload.^18^ Engineered tissues fabricated with polydimethylsiloxane (PDMS) at a 10:1 base-to-crosslinker ratio provided a robust framework for housing H9C2 (rat cardiomyocyte) tissues in a 4 mL media chamber.

H9C2 cells were cultured, transfected, and incorporated into fibrin-based fibers, which were subjected to either static conditions (ST) or pressure overload (PO) (200/10 mmHg) optimized with PE treatment as a positive control (Supplementary Figure 7). Three transfection groups consisting of 20 nM of scrambled siRNA (Scr), commercial calcineurin siRNA (CaN siRNA), and *Cond*-siRNA (2^nd^ Gen) were tested, allowing a comparative analysis of targeted siRNA interventions (Figure 5A).

**Figure 5.**
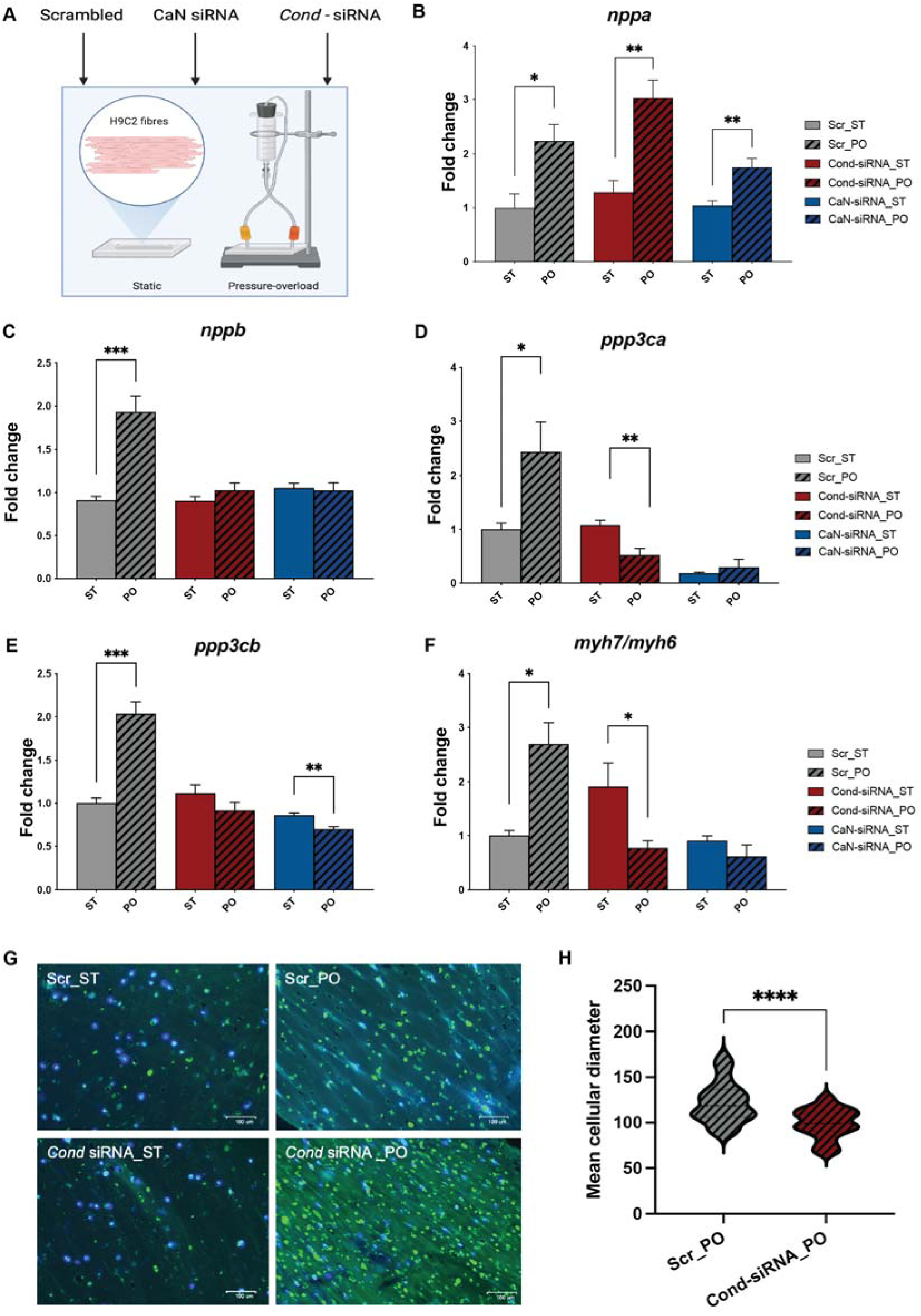
Evaluation of *Cond*-siRNA in the Heart-On-Chip (HOC) Model under HF-like stress conditions. **A.** Schematic representation of the HOC model. H9C2 fibrin-based fibers were subjected to static (ST) or pressure-overload (PO) conditions (200/10 mmHg) following transfection with 20 nM scrambled siRNA (Scr), commercial calcineurin siRNA (CaN siRNA), or Cond-siRNA. Expression levels of stress markers **B.** *nppa,* **C.** *nppb,* **D.** *ppp3ca,* and **E.** *ppp3cb* across experimental groups were estimated using qRT-PCR. **F.** Expression levels of *myh7/myh6* ratio, a marker of cardiac remodeling across all groups assessed by qRT-PCR. Unpaired t-test was performed between ST and PO groups, with significance indicated as *p 0.05, **p 0.01, ***p 0.001. **G.** Wheat Germ Agglutinin (WGA) and DAPI staining of H9C2 fibers under static and pressure-overload conditions. **H.** Quantification of mean cellular diameters Scr_PO and *Cond*-siRNA_PO measured using Image J software. Unpaired t-test was performed between the two groups, with significance indicated as ****p< 0.0001.

Under pressure overload, the expression of *nppa*, a stress marker, was elevated across all treatment groups, indicating a general response to pathological stress (Figure 5B). However, the upregulation of *nppb*, a marker of pathological hypertrophy, was observed only in the Scr_PO group, suggesting that Cond-siRNA mitigated this pathological stress response (Figure 5C). The expression of calcineurin isoforms *ppp3ca* and *ppp3cb* was analyzed to assess the effectiveness of siRNA-mediated knockdown. In the Scr_PO group, both isoforms were significantly upregulated under pressure overload. Treatment with Cond-siRNA selectively reduced the expression of *ppp3ca* under PO conditions while maintaining basal levels under static conditions, demonstrating its conditional activation (Figure 5D). However, Cond-siRNA had no significant effect on *ppp3cb* expression, which remained unchanged across all conditions (Figure 5E). In contrast, CaN siRNA silenced both calcineurin isoforms under all conditions, irrespective of stress, underscoring the specificity of Cond-siRNA for conditional knockdown over the CaN siRNA. The *myh7/myh6* ratio, a well-established marker of pathological cardiac remodeling, was significantly elevated in the Scr_PO group, reflecting hypertrophic changes (Figure 5F). This increase was prevented in the Cond-siRNA_PO group, indicating its efficacy in mitigating pathological remodeling and maintaining a physiological phenotype.

Morphological changes were assessed using Wheat Germ Agglutinin (WGA) and DAPI staining (Figure 5G) with positive staining on mouse heart tissue indicated in Supplementary Figure 7B. Under static conditions (ST), H9C2 fibers appeared compact and uniform. In contrast, exposure to pressure overload (PO) resulted in elongated fibers with wider diameters, reflecting hypertrophic remodeling in the Scr_PO group. Cells in the *Cond*-siRNA_PO group, however, retained a morphology smaller than the Scr_PO group, indicating a protective effect against pressure-overload-induced hypertrophy. Quantification of cellular diameters (Figure 5H) confirmed these observations, with significantly smaller diameters in the *Cond*-siRNA_PO group compared to Scr_PO (p < 0.0001). These findings validate the conditional and stress-specific activation of *Cond*-siRNA, which selectively silences *ppp3ca* under pathological conditions, effectively mitigating hypertrophic remodeling under pressure-overload conditions, highlighting its potential as a targeted therapeutic strategy for HF.

## DISCUSSION

The findings of this study demonstrate the potential of a novel Conditional-siRNA (*Cond*-siRNA) construct in selectively targeting and silencing calcineurin under pathological stress conditions. HF often progresses silently, with patients experiencing subclinical cardiac remodeling long before the onset of symptoms. Early intervention, particularly through targeted therapies, is crucial for preventing irreversible damage and improving long-term outcomes. Recent guidelines highlight the importance of initiating preventive measures in high-risk populations, such as those with hypertension, diabetes, or a family history of HF.^4,5^ Targeted therapies like Cond-siRNA have the potential to fill this critical gap in early intervention. By employing a *nppa*-based sensor system, the *Cond*-siRNA exhibited conditional activation, effectively reducing calcineurin expression and mitigating pathological cardiac remodeling exclusively in pressure-overload conditions. This selective activation addresses a critical limitation of conventional siRNA therapies by ensuring specificity and minimizing off-target effects in non-stressed cardiomyocytes or other cell types.

*In vitro* analysis using Neonatal Rat Ventricular Myocytes (NRVMs) revealed the conditional silencing of calcineurin and downstream inhibition of key signaling pathways, such as NFATc1 nuclear translocation and ERK1/2 phosphorylation. Importantly, the *Cond*-siRNA prevented the upregulation of hypertrophic markers such as β-MHC and reduced cardiomyocyte area under phenylephrine-induced stress without affecting atrophy pathways at relevant concentrations. These observations underscore the efficacy of *Cond*-siRNA in targeting central mediators of pathological hypertrophy while preserving physiological functions in unstressed cells.

The preclinical Heart-On-Chip (HOC) model further validated the stress-specific functionality of *Cond*-siRNA. In this model, the activation of *Cond*-siRNA in response to pressure overload effectively silenced calcineurin isoforms (*ppp3ca* and *ppp3cb*), as evidenced by reductions in their expression levels and normalization of the *myh7/myh6* ratio. Unlike commercial calcineurin siRNAs, which showed silencing in both static and pressure-overload conditions, *Cond*-siRNA demonstrated selective activity, reinforcing its potential for use in dynamic and tissue-specific disease contexts.

The observation that *Cond*-siRNA does not induce muscle atrophy markers, such as *Murf1*, within the therapeutic concentration range highlights its safety and specificity. However, the upregulation of *Murf1* at higher doses of siRNA underscores the need for precise dosage optimization to avoid unintended atrophic effects. This finding aligns with previous studies indicating a dose-dependent threshold for calcineurin silencing^19^ that balances therapeutic efficacy with cellular homeostasis.

Different approaches have been employed to target cardiac hypertrophy, focusing on key signaling pathways and transcriptional regulators that drive maladaptive remodeling. Recent advancements in RNA-based therapies have significantly broadened the therapeutic landscape for cardiac remodeling.^20–22^ Small interfering RNAs (siRNAs) have shown exceptional potential in precisely targeting molecular pathways driving hypertrophy. For example, siRNA-mediated knockdown of mutant *Myh6* transcripts in mouse models has successfully reduced hypertrophic cardiomyopathy, demonstrating the efficacy of gene-specific silencing.^23^ Calcium cycling dysregulation is another critical contributor to cardiac hypertrophy, and targeting pathways regulating this process has proven beneficial. RNA interference strategies targeting phospholamban (PLB) enhance SERCA2a activity, restoring calcium handling and reducing pathological remodeling.^24,25^ Similarly, RNA-based interventions addressing calcium release channels, such as Orai1, have demonstrated protective effects against angiotensin-II-induced pathological remodeling.^26^

Non-coding RNAs, including miRNAs, lncRNAs, and circular RNAs, are emerging as critical regulators and therapeutic targets for various diseases.^27^ miR-30d, for instance, regulates cardiac remodeling through both intracellular signaling and extracellular vesicle-mediated communication. Studies have highlighted its ability to mitigate hypertrophy and fibrosis by modulating critical signaling pathways.^28,29^ Antagonizing miR-208a has shown efficacy in reducing hypertrophy by targeting β-myosin heavy chain (MYH7) expression.^30^ Other lncRNA therapeutic targets against cardiac hypertrophy include H19, whose inhibition reverses pathological remodeling,^31^ Chaer, which modulates chromatin remodeling and epigenetic regulation,^32^ and circular RNAs like circPan3, which regulate hypertrophy through interactions with miR-320-3p.^33^ Among the molecular regulators of remodeling, calcineurin remains a key target due to its role in hypertrophic signaling through the activation of NFAT transcription factors. Its inhibition prevents pathological remodeling and has been validated as an effective therapeutic strategy.^34,35^ Additionally, small molecule inhibitors of calcineurin, have demonstrated improved heart failure outcomes and reduced hypertrophy.^36^ These prior data guided our choice of calcineurin as an initial target in mitigating cardiac hypertrophy. However, calcineurin’s ubiquitous expression across tissues poses a significant challenge, as its inhibition can lead to widespread off-target effects, limiting its clinical application. Our strategy provides a promising alternative by utilizing conditional siRNA, which addresses these limitations through a disease-specific activation mechanism. By leveraging *nppa*, a robust and specific biomarker activated exclusively under pathological conditions, this approach ensures precise therapeutic activation, silencing calcineurin selectively in disease states while sparing normal physiological processes. The conditional siRNA targeting calcineurin has shown promising results in mitigating hypertrophic remodeling in cell culture and preclinical Heart-On-Chip models. Furthermore, the development of cholesterol-conjugated siRNA enhances delivery efficiency, making it a feasible strategy for in vivo applications.

Several challenges remain to be addressed to advance such therapies or expand them into other indications. Most notably ensuring tissue-specific activation, as demonstrated by our use of *nppa* sensors for conditional siRNA activation in cardiomyocytes, for other cell-types or disease states may require considerable work both in determining the appropriate disease-biomarker as well as design of appropriate sensor strands for *Cond*-siRNA. While *nppa* sensors proved effective in this study, their variability highlights the need for expanding RNA sensor repertoires or employing multiplexed strategies to enhance adaptability across diverse pathological scenarios. Robust in vivo validation, including testing in animal models to assess pharmacokinetics, safety, and efficacy under chronic conditions, remains essential. Advanced delivery systems, such as nanoparticles or cardiomyocyte-specific ligands, could improve tissue targeting and minimize degradation, while dosage optimization is crucial to prevent off-target effects like upregulation of muscle atrophy markers. By addressing these challenges, Cond-siRNA constructs hold promise as a personalized therapeutic tool for heart failure, capable of targeting multiple pathological pathways with precision.

In summary, by enabling selective silencing under pathological conditions, conditional siRNA approaches offer an unparalleled combination of specificity and efficacy, minimizing off-target effects while effectively addressing hypertrophy. This methodology bridges critical gaps in current RNA-based therapies and represents a significant advancement in precision medicine for heart failure management. Future studies could expand the Cond-siRNA platform by integrating multiple constructs with distinct sensors and guide RNAs, targeting both hypertrophy and fibrosis to address the multifaceted nature of cardiac remodeling. For example, one Cond-siRNA could silence calcineurin to mitigate hypertrophy, while a second targets pro-fibrotic pathways like TGF-β. This multiplexed approach would enable precise, condition-specific activation, improving therapeutic efficacy by tackling interconnected remodeling mechanisms, paving the way for personalized, multi-target therapies for heart failure while minimizing both off-target effects and undesired on-target effects in other cell-types.

## MATERIALS AND METHODS

### Transverse Aortic Constriction model of non-ischemic HF

Anesthetized male mice aged 8 to 12 weeks underwent a lateral thoracotomy, followed by suture constriction of the transverse aorta against a blunted 25-gaude needle. Control Sham operated mice underwent the same open chest surgery without constriction of the aorta. Heart tissues were harvested 6 weeks post-surgery for RNA isolation. All animal experiments were conducted under approval of the Institutional Animal Care and Use Committee (IACUC).

### Ischemia/Reperfusion (I/R) injury ischemic HF model

I/R was generated by ligating the left anterior descending coronary artery (LAD) using a 7/0 silk thread for 20 minutes, followed by reperfusion, while control Sham mice underwent the same process but without LNA ligation. Mice’s hearts were harvested 6 weeks post-surgery for RNA isolation.

### Annealing and purification of the *Cond-*siRNA

Equimolar concentrations of the sensor, guide and core strands were combined making a total volume of 20µL in 1X PBS. The strands were annealed via thermal annealing (85°C for 3min and 50°C for 1h, followed by cool down). Annealed *Cond-* siRNA constructs were mixed with RNA Loading Dye (component of NEXTFLEX^®^ Small RNA-Seq Kit v3) and loaded in a 10% TBE gel (from Life Technologies EC6275BOX). Gel was run in a XCell *SureLock*™ Mini-Cell (from Life Technologies EI0001) in 1X TBE (BioRad 1610770), 120V for 90minutes. After staining the gel with SYBR® Gold Nucleic Acid Gel Stain (10,000X Concentrate in DMSO) (Thermo Fisher Scientific S-11494), diluted 1x with 1xTBE, for 10minutes, the band corresponding to the *Cond-*siRNA was cut over UV light. The band was pestle (USA Scientific 1415-5390) crushed in 300uL Elution buffer (component of NEXTFLEX^®^ Small RNA-Seq Kit v3) and rotated for at least two hours at RT. The eluted *Cond-*siRNA was filter in a Spin-X tube (Sigma-Aldrich CLS8170-200EA), getting the eluate on the bottom, containing the *Cond-*siRNA.

### Neonatal Rat Ventricular Myocyte (NRVM) isolation

NRVMs were isolated from postnatal day 1 Wistar rat pups using collagenase II and pancreatin based enzymatic digestion, purified via Percoll gradient, and used for experiments 24h post isolation. NRVMs were cultured in DMEM (Life Technologies, Cat.# 11995073) supplemented with 10% horse serum (Thermo Fisher Scientific, Cat.# 26050-088), 5% FBS (Life Technologies, Cat.# 10437028), 1% Penicillin-Streptomycin (Thermo Fisher Scientific, Cat.# 15140122) and 1% L-Glutamine (Thermo Fisher Scientific, Cat.# 25030-081).

### *Cond-*siRNA and commercial siRNA transfection

Transfection of *Cond-*siRNA or other siRNA was performed using Lipofectamine RNAiMax Transfection Reagent (Thermo Fisher Scientific, Cat.# 13778150) into NRVMs in serum free DMEM media (Life Technologies, Cat.# 11995073), supplemented with 1% Penicillin-Streptomycin (Thermo Fisher Scientific, Cat.# 15140122) and 1% L-Glutamine (Thermo Fisher Scientific, Cat.# 25030-081). As positive control of calcineurin silencing a commercial ppp3ca siRNA was utilized (Thermo Fisher Scientific, Cat.# 162268). As negative control AllStars Negative Control siRNA was used (Qiagen, Cat.# 1022076). As positive control of muscle atrophy, miR-29b mimic was used (Thermo Fisher Scientific, Cat.# 4464066) at a 50 pmol concentration.

### RNA isolation and qRT-PCR

For cellular RNA isolation NRVMs, NRCFs and Jurkat T cells, cells were initially lysed with Trizol, and consequently RNA was isolated via chloroform-isopropanol-ethanol protocol. cDNA was then prepared using the High Capacity cDNA Reverse Transcription Kit (Thermo Fisher Scientific, Cat.# 43-688-13), followed by quantitative polymerase chain reaction (qRT-PCR) through the Kapa Sybr Green system for qPCR (Thermo Fisher Scientific, Cat.# KK2601). The sequences of the primers used for quantifying each mRNA are indicated next: *nppa* Forward 5’-GTGCGGTGTCCAACACAGAT-3’, Reverse 5’-TCCAATCCTGTCAATCCTACCC-3’; *nppb* Forward 5’-ACAGCTCTCAAAGGACCAAG-3’, Reverse 5’-GCTTGAACTATGTGCCATCTTG-3’; *myh7* Forward 5’-CCATCTCTGACAACGCCTATC-3’, Reverse 5’-TCTTGGTGTTGACGGTCTTAC-3’; *myh6* Forward 5’-CCATCCTCATCACTGGAGAATC-3’, Reverse 5’-GGTGCCCTTGTTTGCATTAG-3’; *mef2c* Forward 5’-TCTCCGCGTTCTTATCCCAC-3’, Reverse 5’-AGGAGTTGCTACGGAAACCAC-3’; *myocd* Forward 5’-GATGGGCTCTCTCCAGATCAG-3’, Reverse 5’-GGCTGCATCATTCTTGTCACTT-3’; *ddit4* Forward 5’-CCAGTTCGCTCACCCTTC-3’, Reverse 5’-GAAACGATCCCAGAGGCTAG-3’; *ppp3ca* Forward 5’-TTCAGAACGCGTTTATGACGCCT-3’, Reverse 5’-CCTGATGACCTCCTTCCGGG-3’; *ppp3cb* Forward 5’-CCATACTTAGGCGGGAGAAAAC-3’, Reverse 5’-AAGGTATCGTGTATTAGCAGGTG-3’; *murf1* Forward 5’-GAACGACCGAGTTCAGACTATC-3’, Reverse 5’-CCTCCTCCTCCTCTTCAGTAA-3’; *ppp3cb* (for siRNA duplexes 6516 and 6517) Forward 5’-GCTCAAGATGCAGGCTATAGAA-3’, Reverse 5’-CCAACAAATGGTAAAGACCATGTAA-3’; *ppp3cb* (for siRNA duplex 6518) Forward 5’-TGTTGCCTAGTGGAGTGTTG-3’, Reverse 5’-CCGTGGTTCTCAGTGGTATG-3’; *ppp3cb* (for siRNA duplex 6519) Forward 5’-TTAATGTGGAACCTCCCTCACC-3’, Reverse 5’-TTAATGTGGAACCTCCCTCACC-3’; *ppp3cb* (for siRNA duplexes 6520 and 6521) Forward 5’-GAGGATGGATTAGCATGGTC-3’, Reverse 5’-TGGATCTTGTTCAAGTAAGAGT-3’; *actb* Forward 5’-GTGACGTTGACATCCGTAAAGA - 3’, Reverse 5’ - GCCGGACTCATCGTACTCC - 3’. *ppp3ca (human)* Forward 5’-TGGATGTTCTTGCCTCTGAC-3’, Reverse 5’-TGTTTAATCACCATCCCCACC-3’; *nppa (human)* Forward 5’-AACGCAGACCTGATGGATTT-3’, Reverse 5’-TCCTCCCTGGCTGTTATCT-3’; *hprt1 (human)* Forward 5’-TTGCTGACCTGCTGGATTAC-3’, Reverse 5’-CTTGCGACCTTGACCATCTT-3’. mRNA quantification was normalized to the β-Actin (mouse and rat) or HPRT (human) and represented as fold change (2ΔΔ^− Ct^) versus the respective control condition depending on the experimental set up.

### Protein isolation and western blotting

Cytoplasmic and nuclear proteins from NRVMs were extracted through the Thermo Scientific NE PER Nuclear and Cytoplasmic Extraction Kit (Thermo Fisher Scientific, Cat.# 78833). The Pierce™ BCA protein assay (Thermo Fisher Scientific, Cat.# 23227) was performed to quantify lysates’ protein concentration, and 20µg of each sample were used for 4-20% SDS-PAGE electrophoresis. Gels were transferred to PVDF or membranes (BioRad) and blocked with 5%BSA for 1h at RT. Primary antibodies were incubated ON at 4°C rocking at a 1:1000 concentration. The primary antibodies used were the following ones: calcineurin (Cell Signaling Technology, Cat. #2614S), NFATc1 NFATc1(Sigma-Aldrich, Cat. #SAB2101576), Myh7 (Santa Cruz Biotechnology, Cat. # sc-53089), Phospho p44/42 MAPK (Erk1/2) (Thr202/Tyr204) (Cell Signaling Technology, Cat. #4370S) and P44/42 MAPK (Erk1/2) (137F5) (Cell Signaling Technology, Cat. #4695S). Secondary HRP-antibodies (Agilent) were incubated for 1h at RT rocking. Blots were developed using the Supersignal Femto developer (Thermo Scientific, Ct. # 34095).

### Immunofluorescence

NRVMs treated with the *Cond-*siRNA were formalin-fixed for 10min, permeabilized with 0.5% Triton-X100 in PBS, and blocked for 1h at RT. Staining with TroponinT antibody (Thermo Fisher Scientific, Cat. # MA5-12960) (1:250) was performed overnight at 4°C, followed by 2h incubation at RT with the secondary antibody (1:250), and mounted with DAPI containing mounting media (Invitrogen, Cat. # P36935). Images were taken with a Leica SP8 confocal microscope, and cell area quantification was performed through Image J.

### Heart on a chip model

HOC devices were fabricated via soft lithography using polydimethylsiloxane (PDMS) mixed with 10:1 base-to-crosslinker ratio, featuring a bottom flexible membrane, 3 pairs of anchored posts, attached to a ∼1 cm-tall frame via oxygen plasma bonding, to create a chamber that holds 3 engineered tissues and 4mL media. H9C2 cells were cultured until 60–70% confluent, at which point the media in each well of a 6-well plate is replaced with 2 mL OptiMEM. After 2 hours, cells were transfected with 20 nM of siRNA following the manufacturer’s protocol (Lipofectamine). Eight hours post-transfection, the media is switched to DMEM containing 1% P/S (no FBS). Twenty-four hours later, cells are dissociated to create fibers using a mixture of 33 mg/mL fibrinogen (84 µL) and 25 U/mL thrombin (16 µL) per fiber. Once gelled, the fibers are rested in DMEM with 1% P/S and 10% FBS for 48 hours. Following this static culture, the media is changed to serum-free DMEM with 1% P/S for 24 hours prior to experimentation. During the experiment, fibers in the pressure overload group are placed in a 200/10 mmHg loop for 24 hours in serum-free media, while fibers in the static group remain in serum-free media under static conditions.

### Statistics

GraphPad Prism software was used to perform the statistical tests. Comparisons between the two groups were conducted using an independent two-sample t-test. Differences among three or more groups were analyzed using one-way analysis of variance (ANOVA) with Tukey’s post hoc test for pairwise comparisons.

## Author contributions

PG, AMS - designed and performed experiments, analyzed and interpreted the data, wrote the manuscript, and prepared the figures. CG, GL, SH, LS, and RK performed experiments. CA, MS - edited the manuscript. PS, JR, WAG, and SD supplied funding, were involved in the study design, and supervised the work. All authors read and approved the final manuscript.

## Data and materials availability

Will be available upon request

## Supplementary Figures

**Supplemental Figure 1.**
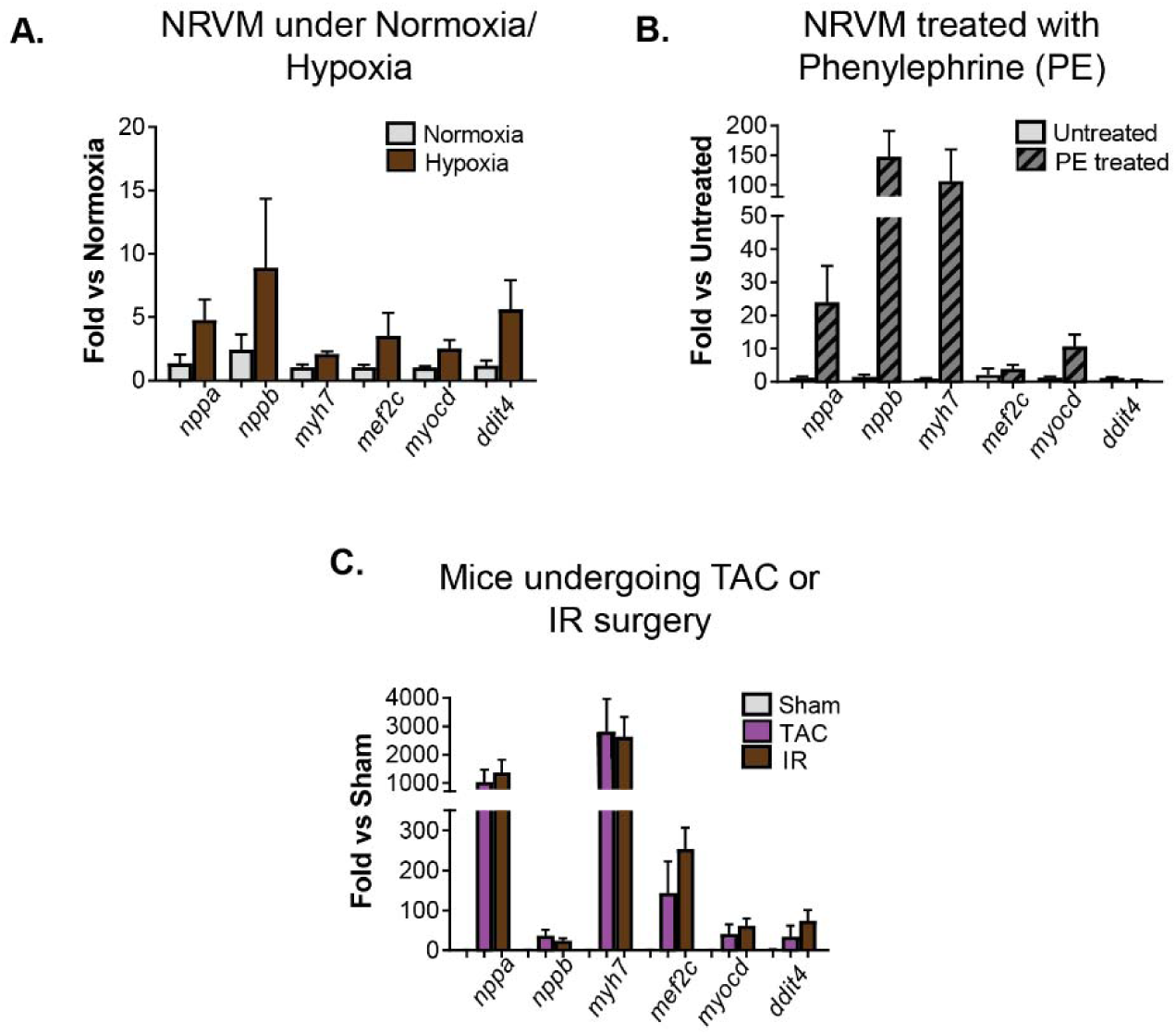
Identification of sensor RNA biomarkers upregulated in response to cardiac stress. Quantification of mRNA levels of cardiomyocyte pathology markers in **A-B**. NRVMs under stress or in mouse hearts **C**. undergoing TAC or IR surgeries. NRVM= Neonatal Rat Ventricular Myocytes, TAC= Transverse Aortic Constriction, IR= Ischemia Reperfusion injury, *nppa*= Atrial Natriuretic Peptide, *nppb*= Brain Natriuretic Peptide, *myh7=* Myosin Heavy Chain beta isoform, *mef2c*= Myocyte Enhancer factor 2C, *myocd*= myocardin, *ddit4*= DNA-damage-inducible transcript 4. Data are shown as Mean ± SEM.

**Supplemental Figure 2.**
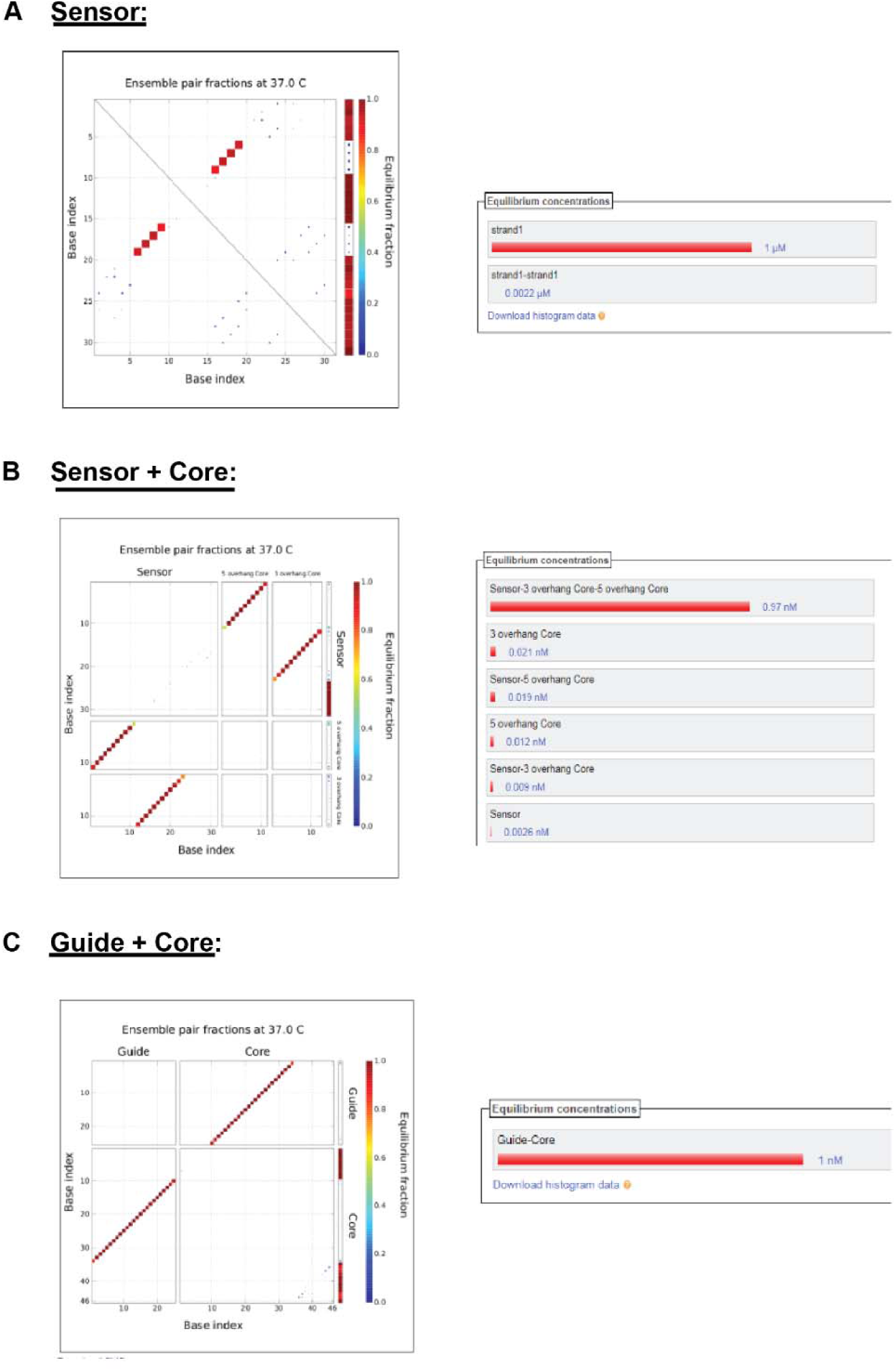
**A-C.** Nupack software’s plots ranking the thermodynamic stability of the complexes formed between the different strands of the *Cond-*siRNA.

**Supplemental Figure 3.**
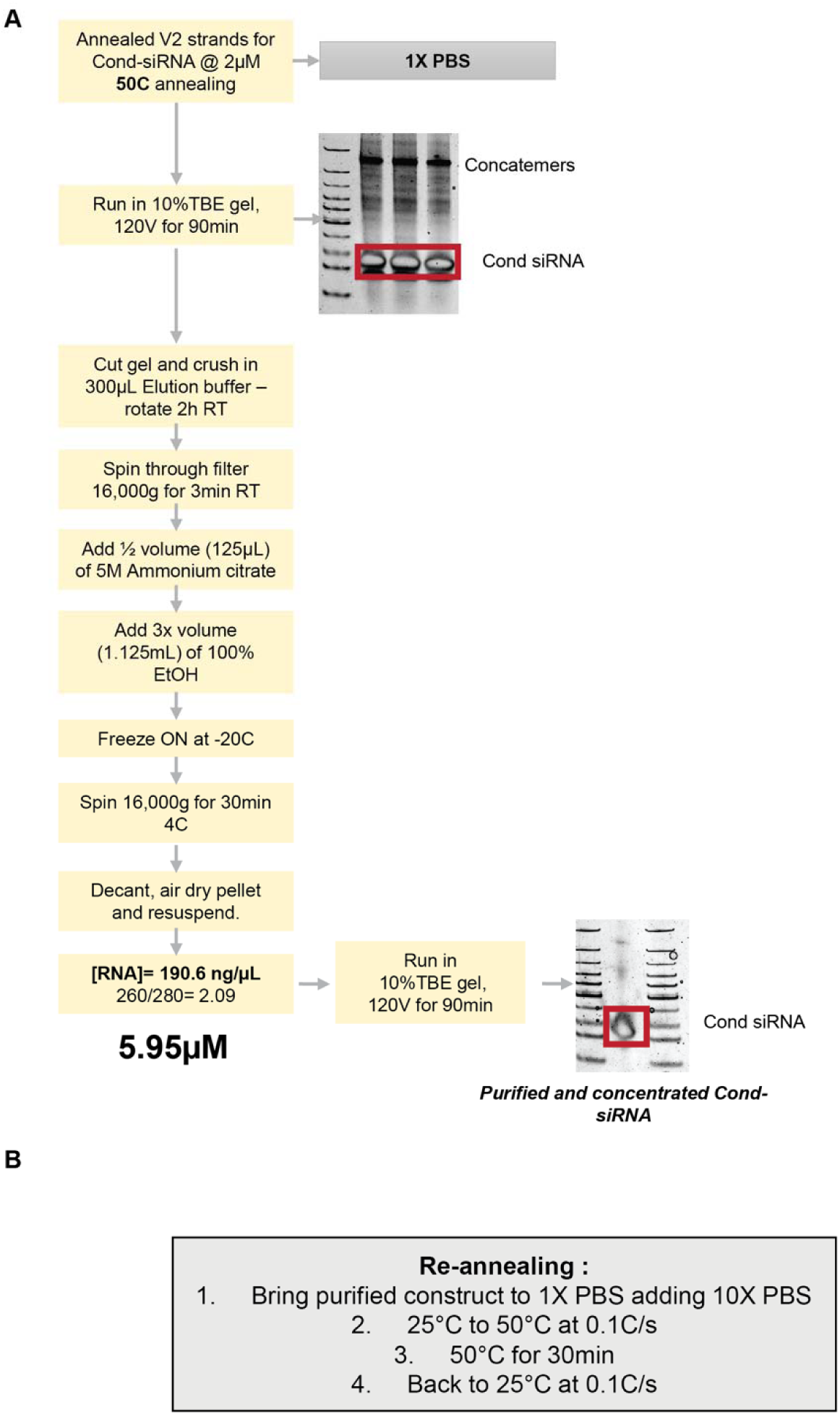
**A.** Protocol for strand annealing at high concentration, followed by purification to obtain the well-assembled Cond-siRNA and subsequent concentration. **B.** Protocol for re-annealing.

**Supplemental Figure 4.**
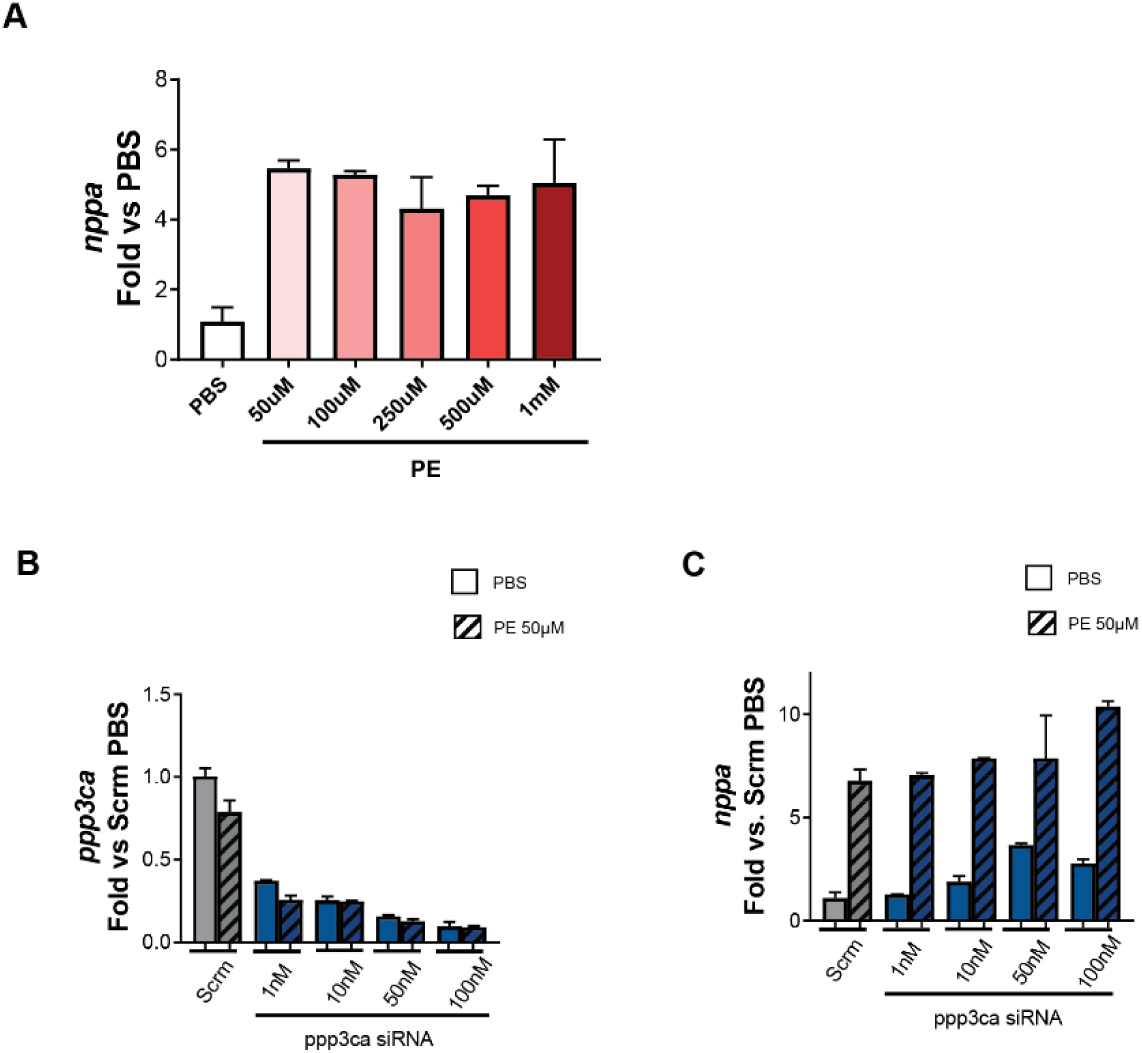
**A.** *nppa* expression levels in NRVM treated with increasing concentrations of PE for 48 hrs. Data is represented as Fold vs PBS. **B.** Calcineurin *ppp3ca* isoform and **C.** *nppa* mRNA expression in NRVMs transfected with increasing concentrations of commercial siRNA targeting *ppp3ca*, and treated with 50µM PE. Data is represented as fold vs Scramble PBS.

**Supplemental Figure 5.**
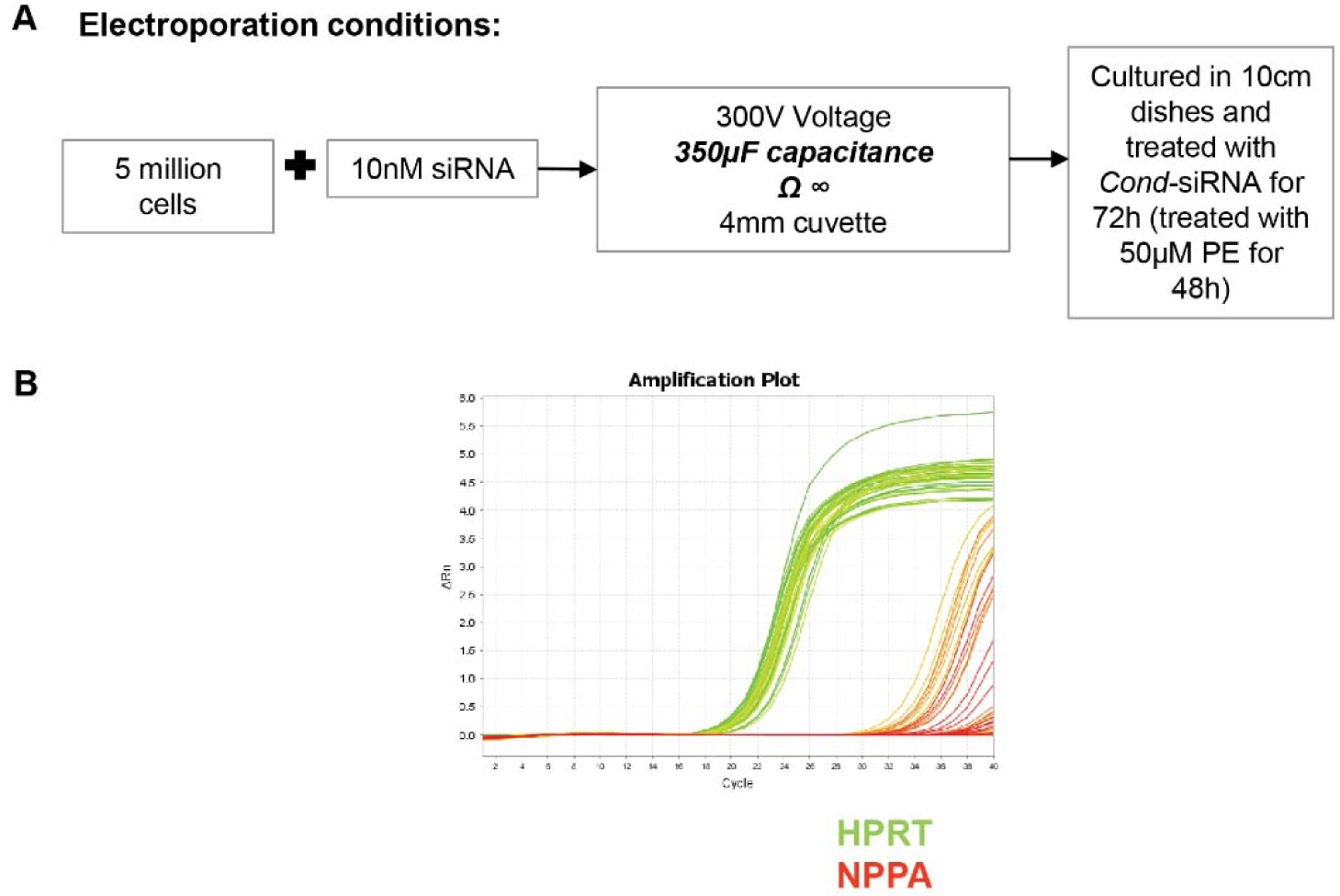
**A**. Flowchart of the Jurkat cell electroporation and culture conditions. **B**. Representative amplification plot of *nppa* and *hprt* (house-keeping gene) in Jurkat cell qRT-PCR.

**Supplemental Figure 6.**
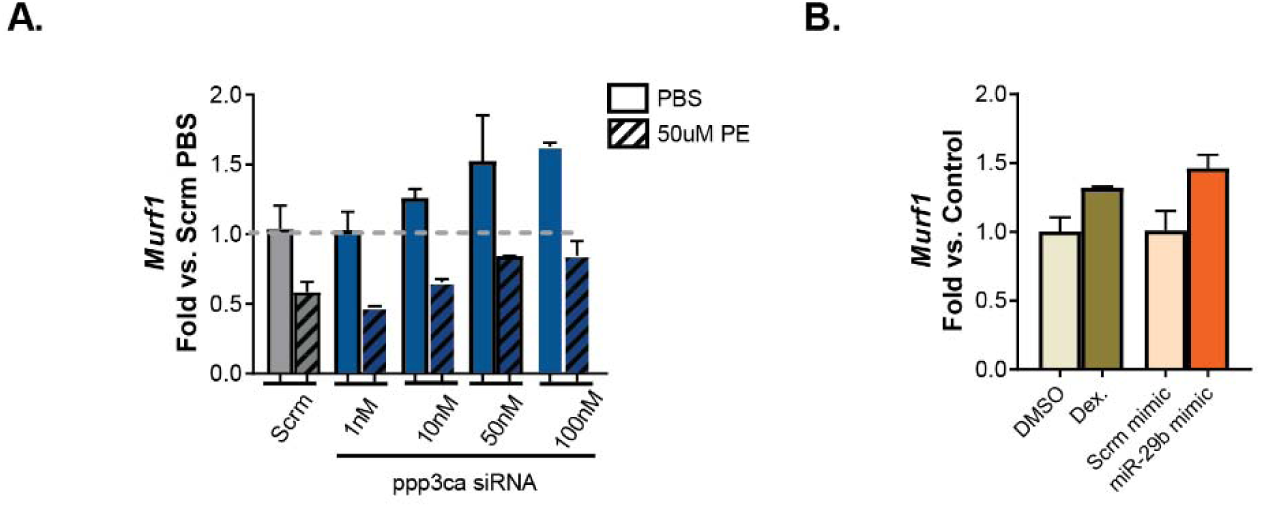
**A.** *Murf1* expression levels in NRVM treated with increasing concentrations of commercial siRNA and PE. Data is represented as fold vs PBS. **B.** *Murf1* expression levels in NRVM treated with 50µM dexamethasone for 24h or transfected with 50pmol miR-29b mimic for 48h. Data is represented as fold vs its respective control.

**Supplementary Figure 7.**
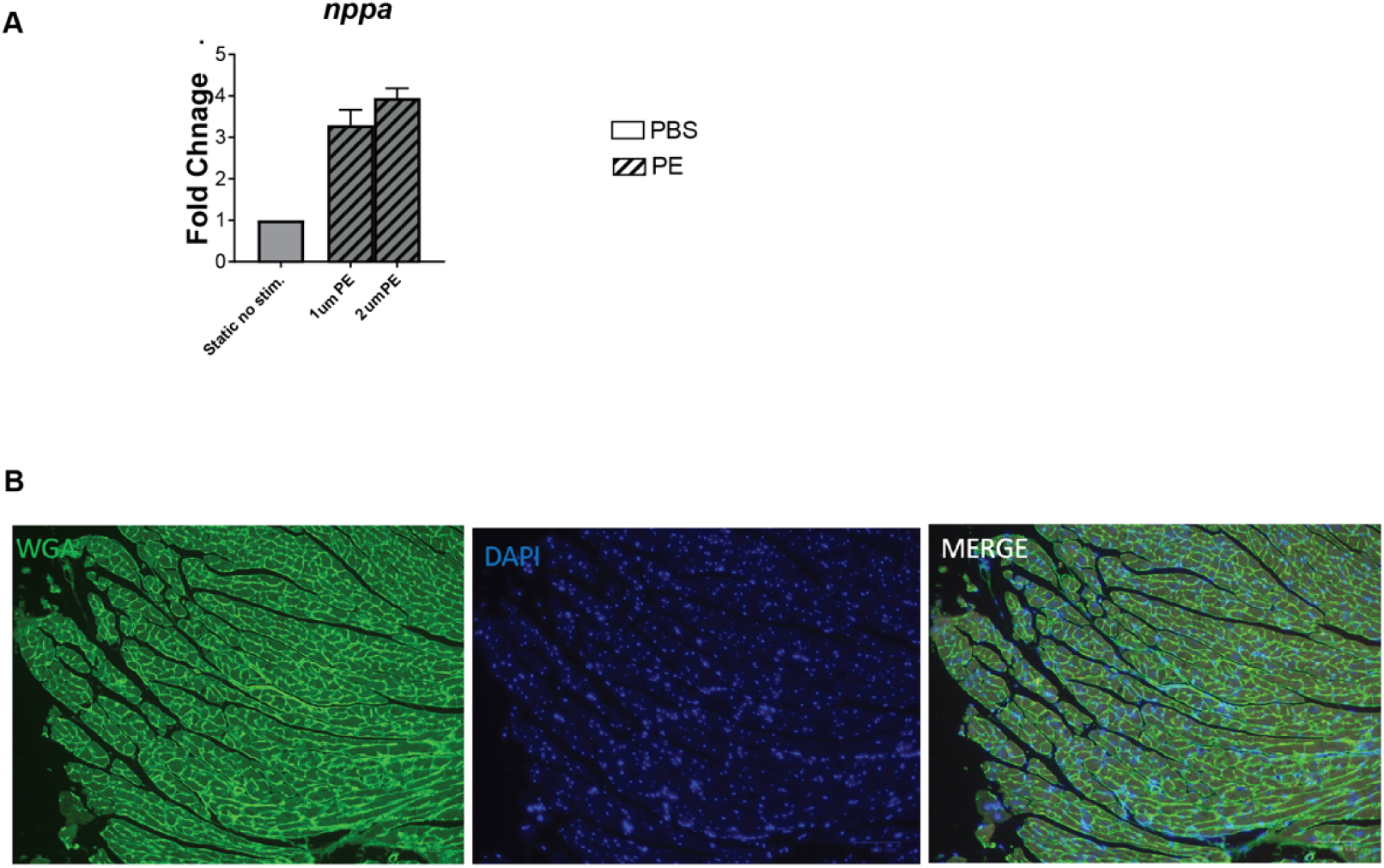
**A.** Optimal stimulation condition for inducing ANP upregulation on the organ-on-a-chip model upon PE treatment evaluated using qRT-PCR. **B.** Positive immunofluorescence staining of Wheat Germ Agglutinin (WGA, Green) and DAPi nuclear staining (Blue) on heart tissue.

